# Mistargeted retinal axons induce a synaptically independent subcircuit in the visual thalamus of albino mice

**DOI:** 10.1101/2024.07.15.603571

**Authors:** Sean McCracken, Liam McCoy, Ziyi Hu, Julie Hodges, Katia Valkova, Philip R. Williams, Josh Morgan

## Abstract

In albino mice and EphB1 knock out mice, mistargeted retinal ganglion cell (RGC) axons form dense islands of axon terminals in the dorsal lateral geniculate nuclei (dLGN). The formation of these islands of retinal input depends on developmental patterns of spontaneous retinal activity. We reconstructed the microcircuitry of the activity dependent islands and found that the boundaries of the island represent a remarkably strong segregation within retinogeniculate connectivity. We conclude that, when sets of retinal input are established in the wrong part of the dLGN, the developing circuitry responds by forming a synaptically isolated subcircuit within the otherwise fully connected network. The fact that there is a developmental starting condition that can induce a synaptically segregated microcircuit has important implications for our understanding of the organization of visual circuits and for our understanding of the implementation of activity dependent development.

## INTRODUCTION

Activity dependent synaptic remodeling is a process in which patterns of neuron depolarization and synaptic transmission determine whether a given synapse will be maintained, enhanced, or eliminated (Katz and Shatz 1996; J. W. Lichtman and Colman 2000; Redfern 1970; Jeff W. Lichtman 1977; Crepel, Mariani, and Delhaye-Bouchaud 1976). This process is important for both the formation of circuits during development and for the creation of long-term memory. However, it has proven difficult to perform experiments in which specific patterns of neural activity can be mapped onto features of the synaptic organization of circuits. To understand how experience is shapes the cellular organization of circuits, we look to a model system in which stereotyped patterns of activity generate a stereotyped functional architecture.

In amniotes (birds, reptiles, mammals), the dorsal lateral geniculate nucleus (dLGN) provides the most direct link between the retina and visual cortex. The dLGN’s functional architecture is similar to that of their immediately upstream circuit, the innerplexiform layer of the retina. Both neuropils consist of multiple topographic maps of visual space stacked together as functionally distinct layers (Reese 1988). Unlike the architecture of the retina, the functional architecture of the dLGN depends heavily on an extended period of activity dependent synaptic remodeling (Chen and Regehr 2000; Stellwagen and Shatz 2002; Lee, Eglen, and Wong 2002; Godement, Salaün, and Imbert 1984; Hahm, Langdon, and Sur 1991). Blocking the spontaneous waves of activity disrupts the refinement of dLGN retinotopy (Grubb et al. 2003; Pfeiffenberger, Yamada, and Feldheim 2006) and the normal segregation of left and right eye inputs (Feller 2009).

The innervation of the dLGN by the retina starts with molecular gradients guiding axons to roughly the correct location (Sitko, Kuwajima, and Mason 2018) where they form synapses with many more thalamocortical cells (TCs) than they will innervate in the adult (Chen and Regehr 2000). The retina than generates spontaneous waves of activity in which RGCs with similar positions and receptive field properties fire together (Meister et al. 1991). The temporal correlation of this RGC firing is thought to help refine the innervation of the dLGN according to Hebbian rules (Butts, Kanold, and Shatz 2007; Lee, Eglen, and Wong 2002; Hebb 1949). By selecting a set of RGC inputs that fire at the same time, TCs can ensure that their inputs are driven by a single region of space and have similar functional properties. The result is a refined map of visual space where nearby TC share functional properties and, often, RGC inputs (J. L. Morgan et al. 2016).

What would happen if a small set of RGC axons took a wrong turn early in development and found themselves surrounded by RGCs axons with very different activity patterns? The Hebbian model would predict that these inputs might not be integrated into the surrounding network. Rather, they might capture a set of TCs that is exclusive to the mistargeted RGCs. The result would be a synaptically segregated retinogeniculate subcircuit, an organization not normally observed in mouse dLGN (J. L. Morgan et al. 2016).

Examination of the RGC boutons distributions in mice in which the initial targeting of RGCs to the dLGN is disrupted reveal structures that consistent with the predicted segregated microcircuit. In albino mice (B6(Cg)-Tyr^c-2J^/J,) (Rebsam, Bhansali, and Mason 2012) and EphB1-knock out mice (Rebsam, Petros, and Mason 2009), sets of RGC axons that would normally project to the ipsilateral dLGN, instead project to the contralateral dLGN. Some of these mistargeted axons form islands of intense bouton labeling within the dLGN. Blocking developmental retinal waves, prevents the formation of these islands (Rebsam, Petros, and Mason 2009; Rebsam, Bhansali, and Mason 2012). These islands of RGC boutons are, therefore, strong candidates for being the segregated microcircuits predicted by a Hebbian model of the results of axon mistargeting.

What was unknown, based on these previous studies, was the microcircuitry underlying the islands of dense RGC bouton labeling. Here, we use correlated light to electron microscopy (CLEM) to test whether the islands of boutons in the albino mouse dLGN form synapses or are simply a tangle of axon terminals that could not find a target. We find that RGC axons in the island form the complex retinogeniculate synaptic glomeruli typical of normal dLGN. We then test the Hebbian prediction that TCs should be innervated by either the island or non-island RGC boutons, but not by both. We found that the island does, in fact, reflect a segregated microcircuit where mistargeted axons capture an exclusive set of TCs. Finally, we examined the extent to which the retinogeniculate circuit segregation extends to other cell types in the albino dLGN. Our results demonstrate, not only that the activity-dependent RGC island constitutes a segregated microcircuit, but that the segregation has a significant impact on the cellular organization of the dLGN.

## METHODS

### Animals

Albino mice (B6(Cg)-Tyr^c-2j^/J) and GAD67-GFP (CB6-Tg(Gad1-EGFP)G42Zjh/J) mice were obtained from Jackson labs. All mice were between two and three months old. Optical examination of dLGNs included both male and female mice. The mouse on which vEM was performed, was male. All procedures in this study were approved by the Institutional Animal Care and Use Committee of Washington University School of Medicine (Protocol # 23-0116) and complied with the National Institutes of Health Guide for the Care and Use of Laboratory Animals.

To obtain tissue, mice were transcardially perfused with either 4% paraformaldehyde in 0.1 M phosphate buffered saline (PBS)(optical imaging only) or 2% paraformaldehyde and 1% glutaraldehyde in 0.1 M PBS. Brains were removed and post fixed for 30 minutes to an hour. Brains were then cut to 50 to 200 μm thick coronal sections using a Compresstome (Precisionary Instruments) vibratome.

### Optical labeling

RGC axons were labeled by intraocular injection of choleratoxin-B (CtB) bound to either Alexa-488, Alexa-555, or Alexa-647. Cell nuclei were labeled by incubating vibratome sections in DAPI for 20 minutes or by mounting in Fluoromount with DAPI (ThermoFisher Scientific). Vibratome slices were partially cleared by soaking in 2,2’-thiodiethanol (TDE) or by repeated applications and removal of Vectashield (Vector Laboratories).

### Optical imaging

Brain slices were first imaged using a widefield epifluorescence microscope (Leica). Slices were then imaged using an FV-1000 confocal microscope (Olympus). Scans typically included two channels of CtB fluorescence acquisition, a DAPI channel, a reflected light channel (no laser filtering), and an auto- fluorescence channel (excitation wavelength for aldehyde and without corresponding label). Images of GAD67-GFP labeled LINs were acquired with a Zeiss LSM 800 (Zeiss) confocal microscope in Airyscan mode. Images were acquired with low laser, maximum scan speed, and averaging between 3 and 8 scans. High-resolution images were acquired with a 60x 1.4 NA oil objective. For presentation, the brightness, contrast, and gamma of each channel has been independently adjusted. The goal of this adjustment is to make both the tissue background signal and cellular structures visible without saturating pixels.

### Electron microscopy

After optical imaging of the full vibratome brain slice, tissue was removed from the slide and refixed with 2% paraformaldehyde and 2% glutaraldehyde. The dLGNs were then isolated from the rest of the brain slice and washed with 0.1 M Cacodylate buffer. Tissue then treated with osmium tetroxide -> ferrocyanide -> thiocarbohydrazide -> osmium tetroxide -> uranyl acetate -> lead aspartate (Hua, Laserstein, and Helmstaedter 2015; J. L. Morgan et al. 2016). Forty-nanometer-thick ultrathin sections were collected using an automatic tape collecting microtome (RMC ATUM, Leica UC7 ultramicrotome, (Schalek et al. 2012)). Sections were collected onto carbon coated Kapton (gift from Jeff Lichtman lab) and then mounted onto silicon wafers.

Wafers were mapped on a Zeiss Merlin scanning electron microscope using WaferMapper (Hayworth et al. 2014). Mapping includes acquiring a low-resolution image of each tissue section. The resulting image volume was compared to optical images of the tissue section to target further imaging. Before high resolution images were acquired, sections were plasma etched with an ibss plasma asher (ibss Group, Inc.) to enhance contrast and surface stability (J. L. Morgan et al. 2016). High-resolution (2 nm pixel) images were acquired from targeted regions of the island and exclusion zone. The vEM volume used for most of the circuit reconstruction was acquired at 20 nm pixel size. Both types of images were acquired with the in-lens secondary electron detector at 1kev landing voltage and 1-3 nA. The 3D image volume was aligned by ScalableMinds.

### vEM rendering and analysis

Circuit reconstruction was performed via manual tracing in VAST (Berger, Seung, and Lichtman 2018). Segmentations were imported from VAST to CellNav (https://github.com/MorganLabShare/CellNav) for analysis and visualization. 3D renderings and analysis can be regenerated by downloading CellNav version AlbinoIsland and the AlbinoIsland CellNav cell library (https://sites.wustl.edu/morganlab/data-share/). On executing ‘runCellNav’, select the AlbinoIsland CellNav directory. All cells and synapses included in this study are available to view. The rendering of the island and non-island boutons can be generated by Menu-> RunScripts -> Publication -> AlbinoIsland -> jm_KxS_showGreenMagentaBoutons.m. The images used in Supplemental Figure 2 can then be generated by running Menu -> RunScripts -> Publication -> AlbinoIsland -> jm_KxS_showCellsAndRGC_snapShots.m. The plots in Figure 2 can be generated by running Menu -> RunScripts -> Publication-> AlbinoIsland -> PlotCBtoSynVector. Synapses associated with particular cells can be rendered by selecting a cell or cell type from the Select Cells panel and then clicking ‘Cell’ or ‘Type’ in the Groups panel. The cell group can then be defined as the pre or post synaptic cell in the ‘Synapses’ tab. Synapses can be filtered by pre or post synaptic cell type use the ‘preClass’ or ‘postClass’ drop down menus.

### Monte Carlo analysis

We performed a Monte Carlo analysis to gauge the extent to which the connectivity of the reconstructed circuit deviates from a circuit in which there is no selection against a TC being innervated by both island and non-island RGC boutons. We selected 9 reconstructed TCs whose cell bodies were determined to be in the exclusion zone. For these TCs, we counted how many primary dendrites were innervated by RGC boutons or whose downstream dendrites were innervated by RGC boutons. We then ran a model 100,000 times in which each primary dendrite had an equal probability to be connected to the island or non-island RGC boutons. We then counted how many TCs in each trial were innervated by only the island boutons or only the non-island boutons. We reported the value (4 TCs) for which 99% of the Monte Carlo results produced equal of fewer captured TCs.

### Statistical design

In this project, we have tested the hypothesis that the mistargeted RGC axons in albino mice form a synaptically segregated retinogeniculate circuit. This hypothesis was qualitative. Prior to performing this study, we did not have a reasonable basis for formulating a quantitative hypothesis regarding how much synaptic segregation would constitute a segregated microcircuit. We, therefore, did not design a null hypothesis significance test to reject the null hypothesis that the circuit was not segregated.

We did compare our observed circuit reconstruction to a model of a circuit without segregation and found that our EM reconstruction demonstrated a substantial deviation from the no-segregation prediction. This model is based on 9 cells from the same piece of tissue. There is no control for variation between individuals. None the less, the test demonstrates that it is possible for a mouse dLGN to produce a highly segregated retinogeniculate subcircuit. We also report that the optically identifiable correlate to the structures we describe in our EM reconstruction are present in 12 of 12 albino mice examined and match the pattern reported in the previous literature (Rebsam, Bhansali, and Mason 2012).

## RESULTS

### The albino island is surrounded by an RGC bouton exclusion zone

We first replicated previous studies of the distribution of RGC boutons in the albino mice. We labeled the RGCs of 12 albino mice with the anterograde neuronal tracer choleratoxin-B (CtB) and examined the dLGNs with widefield epifluorescence imaging of serial vibratome sections. CtB labeling is particularly bright within synaptic boutons and injecting each eye with a different color of fluorescently bound CtB (Alexa 488, 568, 647) is a standard method of revealing retinogeniculate synaptic connectivity.

Consistent with previous studies, we found that the dLGNs of all 12 mice examined showed bright, distinct islands of contralaterally projecting RGC boutons (Figure 1).

**Figure 1:**
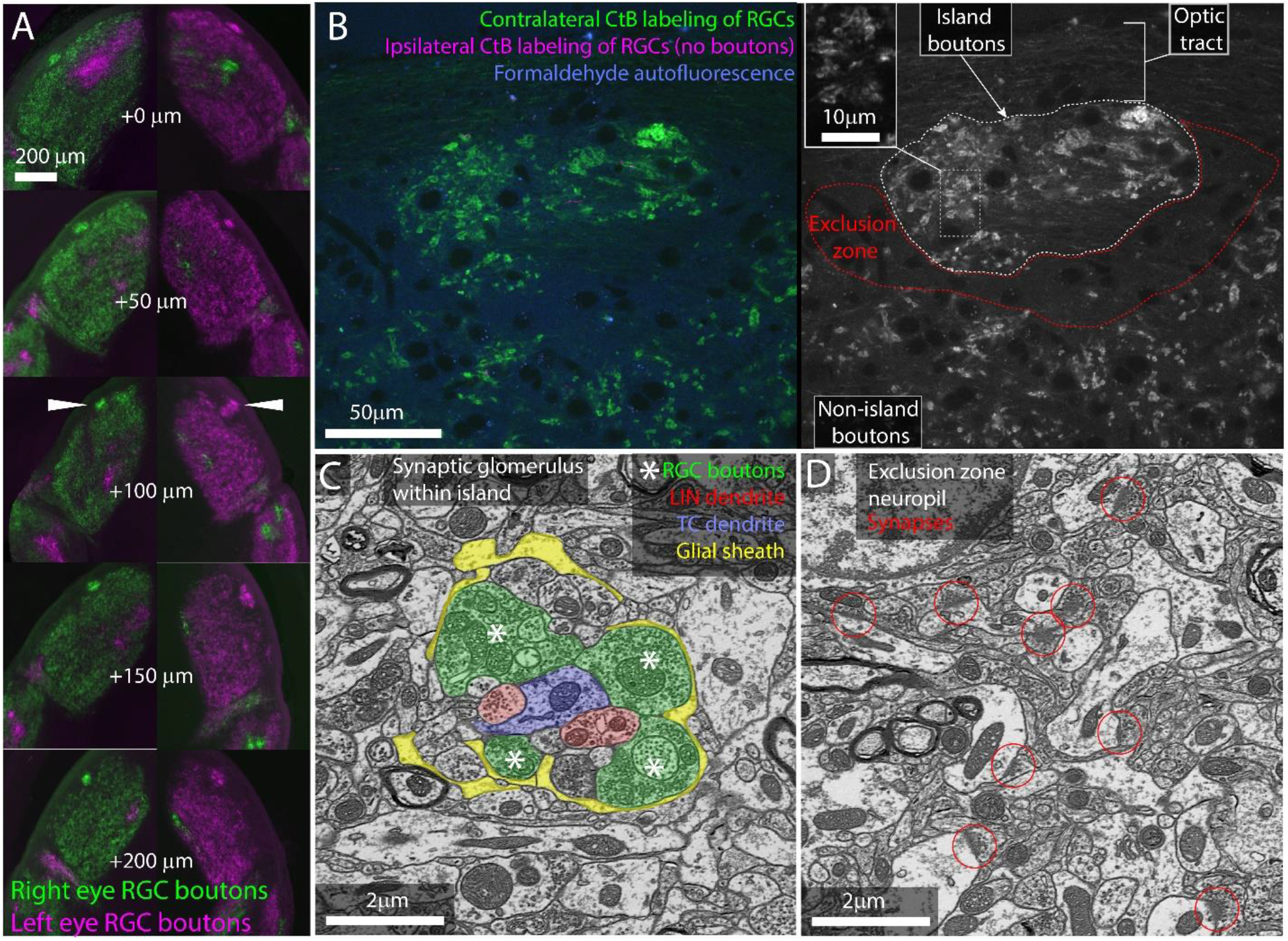
The island of RGC boutons represents a segregation and concentration of a subset of otherwise normal retinogeniculate connections. A) Serial vibratome sections through the left and right dLGNs of an albino mouse. From top to bottom section progress rostral to caudal. CtB labeling of RGCs from the left (magenta) and right (green) eyes. White arrowheads indicate island. B) Single confocal section through CtB labeled albino island. Left shows background fluorescence (blue), contralateral RGC boutons (green) and an absence of ipsilateral axons (magenta). Right shows CtB labeled RGC boutons from contralateral eye only. Inset shows enlarged view of RGC boutons. C) Electron micrograph of retinogeniculate glomerulus in the albino island. RGC boutons are indicated by asterisks. D) Electron micrograph of non-glomerular neuropil of the exclusion zone. Red circles indicate non-glomerular synapses. Both the synaptic glomerulus in the island and the feedback synapses in the exclusion zone appear ultrastructurally normal.

**Figure 2.**
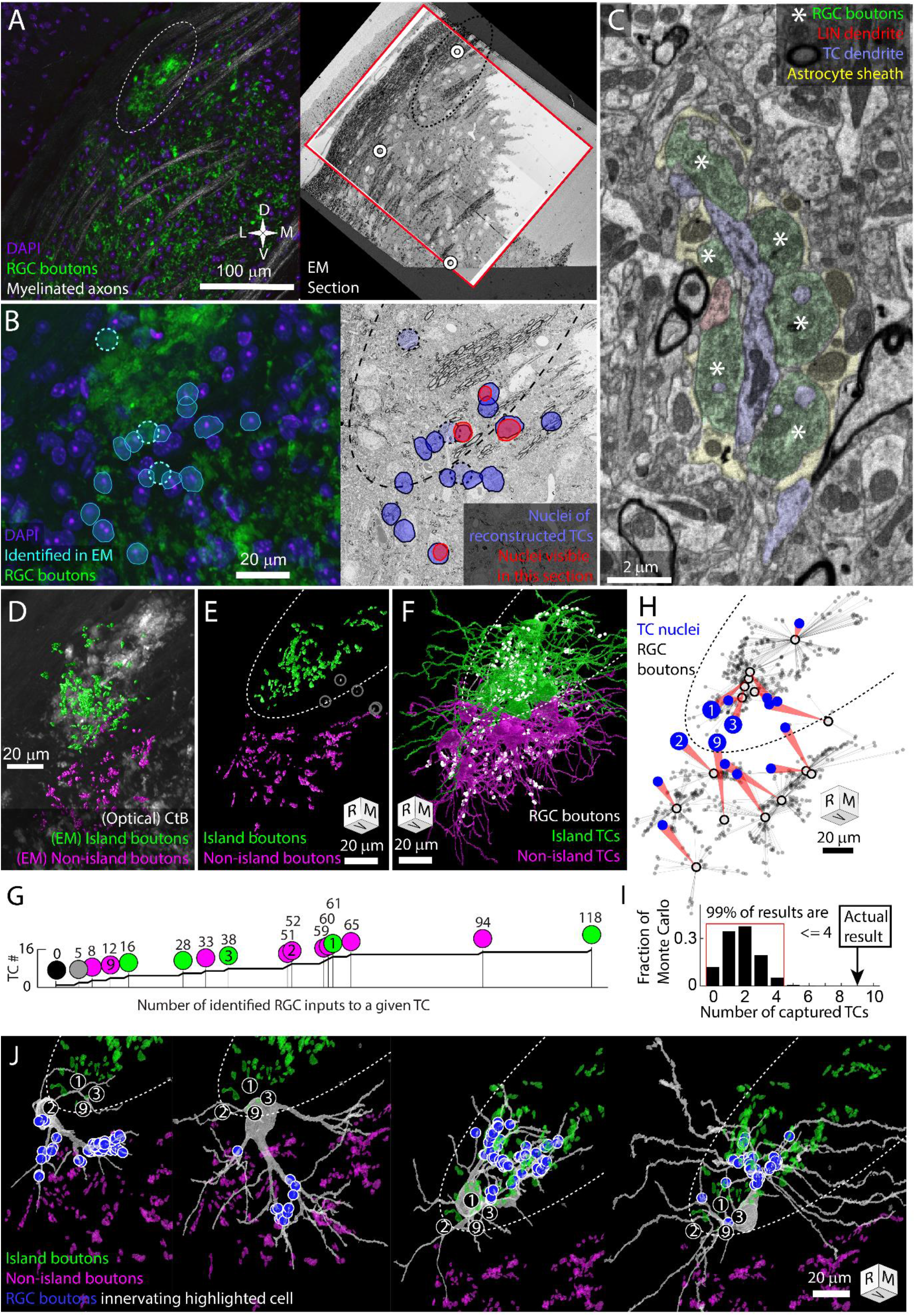
The RGC island represents a segregation of retinogeniculate synaptic connectivity. Thalamocortical cells are either innervated by island RGC boutons or non-island RGC boutons. Dotted oval indicates the boundary of the RGC island in multiple panels. Four cells (TC1, TC2, TC3, TC9) are identified in panel B, G, H, and I. Tissue orientation is indicated by compass rose or rotation cube (Rostral/Medial/Ventral). A) Matching of optical images and EM section overview. Red rectangles indicate the boundaries of the EM volume used to trace TCs. White circles indicate blood vessels matched between optical images and EM images. B) Closer view of light and EM images showing nuclei of EM-reconstructed TCs. In the optical image (left), DAPI labeled nuclei (blue) of reconstructed TCs are outlined (cyan) relative to RGC boutons (green). The position of reconstructed TC that are not visible in the confocal image stack are indicated by circles. The nuclei labels are mapped onto the EM section (right). The nuclei of reconstructed TCs visible in the EM section are highlighted in red. C) Example of electron micrograph with 20 nm pixel size in which TC dendrite and associated glomerulus are annotated. RGC boutons are indicated by asterisks. D) EM reconstructed island (green) and non-island (magenta) RGC boutons that were found innervating reconstructed TCs. EM boutons are overlayed on corresponding optical image of CtB labeled RGC boutons. E) Rotation of EM labeled RGC boutons to show clearest separation between island (green) and non-island boutons (magenta). Boutons innervating the one TC without a clear island or non-island identity are highlighted in white. F) Partial reconstructions of TCs indicated in panel B. TCs are color coded by whether the receive their input from the island or non-island RGC boutons. Positions of RGC inputs are indicated by white dots. G) Cumulative curve of the number of RGC boutons innervating reconstructed TCs. Circles indicate island / non-island identity. Numbers above the circles are the number of RGC boutons innervating the TC. Number within the circle indicates TC IDs used in other panels. H) Plot of the relationship between TC cell body (blue circle) and the location of the RGC inputs that innervate it (grey dots). Average synapse position is indicated by black circle. I) Results of Monte Carlo simulation predicting how many TCs would be innervated only by island or only by non-island RGC boutons if TC dendrites are independent of one another. Red bracket encloses 99% of results. J) EM reconstructions of the four example TCs highlighted in previous panels. Circled numbers indicate the relative position of the four example cells. The location of RGC inputs are indicated by blue circles. Island and non-island RGC boutons innervating other TCs shown in green and magenta. TC cells with soma in the exclusion zone have asymmetric dendritic arbors that reflect their exclusive connectivity to either the island or non-island RGC boutons.

We next examined the cellular organization of the albino island in the same samples using confocal microscopy. Confocal imaging of the CTB labeled albino dLGNs showed that CtB labeling of RGC boutons within the island resembled labeling of RGC boutons outside of the island with the exception that the density of labeling was higher within the island (Fig 1B). Both inside and outside of the island, we observed the fluorescent profile of bright dots surrounding a dark spot. This profile is consistent with synaptic glomeruli in which labeled RGC boutons clustered around unlabeled TC dendrites (Fig 1B,C).

Confocal imaging also made it clear that the island was surrounded by an RGC-bouton-free region we term the exclusion zone. Examination of this exclusion zone with autofluorescence, reflected light, and immunolabeling for synaptophysin (Figure 1B, Supplementary Figure 1) demonstrates that the exclusion zone was composed of a mix of neuropil, cell nuclei, myelinated fibers, and an occasional blood vessel. We did not detect an anatomical barrier that would preclude RGC boutons from forming in this region.

We next asked whether the optically labeled island boutons formed normal synaptic connections. We used correlated light and electron microscopy (Friedrichsen et al. 2022) to examine the ultrastructure of one albino island. We found that RGC boutons in the island participated in normal retinogeniculate synaptic glomeruli (Fig 1C). RGC boutons were identified by their light mitochondria, large size, and large synaptic vesicles (Hamos et al. 1987). Thalamocortical dendrites were distinguished from local inhibitory neuron dendrites by the presence of spines and the absence of synaptic outputs. RGC boutons were found in clusters surrounding TC dendrites. The RGC boutons form normal synaptic triads innervating local inhibitory neuron boutons that then form inhibitory synapses onto thalamocortical cells. As in normal dLGN, the cluster of RGC and local inhibitory neuron boutons are encapsulated by a glial sheath (Pecci Saavedra and Vaccarezza 1968).

Consistent with the CtB labeling, EM of the exclusion zone revealed large regions of dLGN neuropil with no RGC boutons (Figure 1D). Aside from the lack of RGC glomeruli, the exclusion zone appeared healthy. Normal mouse dLGN neuropil can be divided into RGC-bouton-associated glomerular neuropil (Figure 1C, colored) and non-glomerular neuropil. The non-glomerular neuropil consists primarily of the distal dendrites of TCs and their inputs from the visual cortex and thalamic reticular nucleus (Sherman and Guillery 1996). The synaptic neuropil of the exclusion zone was composed of non-glomerular synapses and their associated neurites (Figure 1D).

Our examination of RGC boutons in the dLGN of the albino mouse, therefore, rules out the possibility that the islands are dead-end neuromas buried in the optic tract. Rather, the bright islands of CtB labeling observed in the albino dLGN represent a high concentration of ultrastructurally normal RGC boutons embedded in healthy dLGN neuropil.

### An exclusive set of thalamocortical cells are captured by the island RGC boutons

Does the spatial segregation of RGC boutons reflect a segregation in the functional connectivity of the albino dLGN? TCs are the sole output neuron of the dLGN and their receptive field properties are largely the product of which RGCs innervate them (Usrey, Reppas, and Reid 1998). The pattern of RGC innervation of thalamocortical cells, therefore, defines the functional architecture of the dLGN. In previous EM reconstructions of mouse dLGN circuitry, we found that nearby thalamocortical cells tend to be innervated by the same RGC axons (J. L. Morgan et al. 2016). By contrast, If the exclusion zone around the RGC island represents a segregation of retinogeniculate circuitry, we would expect to find a hard boundary in connectivity. TCs near the exclusion zone would be innervated by either the island or non-island RGC boutons, not both.

To test this hypothesis, we collected a serial section EM volume to reconstruct the arbors and synaptic connectivity of thalamocortical cells near the exclusion zone. The EM volume was obtained from an optically characterized albino dLGN and consisted of 2631 40 nm thick coronal sections (Figure 2A,B). From each section, we acquired a 220 μm x 220 μm image mosaic that encompassed most of the albino island, the exclusion zone, and the surrounding non-island dLGN neuropil (105 μm of rostro-caudal z-depth). Images were acquired with a 20 nm pixel size. For analysis, the voxel size was treated as 26 nm x 20 nm x 40 nm to compensate for tissue compression during cutting. The large pixel size (usually 4 nm for circuit reconstruction) allowed for rapid acquisition of a large vEM volume while still allowing for the tracing of large features such as TC dendrites and RGC boutons (Figure 2C).

We traced 15 thalamocortical cells whose nuclei were distributed across the RGC island exclusion zone (Figure 2B). TC dendrites were traced until they left the volume or until they had clearly transitioned into a distal dendrite morphology. All reconstructions should be considered partial. The transition from proximal to distal dendrite morphology was identified as the point where dendrites become thin and unbranched and form spines that are innervated by small cortical feedback boutons as opposed to RGC boutons. After tracing the TCs, we labeled the RGC boutons innervating the TCs.

The RGC inputs identified in the EM reconstruction of TCs covered approximately a third of the island visible in the confocal imaging of the same tissue (Figure 2D). 3D rotation of the reconstructed boutons revealed a clear exclusion zone between the reconstructed island boutons and the surrounding boutons (Figure 2E). The exceptions were RGC boutons associated with TC17. This TC was innervated by only five RGC boutons, three of which were in the middle of the exclusion zone (Figure 2E gray circles).

We found that 14 TCs were innervated either exclusively by the island (6 TCs) or exclusively by RGC boutons outside of the island (8 TCs, Figure 2F-H). There were no mixed non-island / island TCs. There was one TC (mentioned above) that received inputs in the exclusion zone and one TC on which no RGC inputs could be found. The wide range of the number of RGC boutons innervating TCs (0-118, Figure 2G) may reflect the fact that, in the dorsal mouse dLGN, synaptic inputs from the superior colliculus can take the place of RGCs as a primary driving synapse (Bickford et al. 2015).

We next compared our observed results for the rate of complete capture of a TC, to what we would expect if there was no preference for being innervated by only island or only non-island RGC boutons. We designed a simple Monte Carlo simulation based on the 9 TCs whose cell bodies were most clearly in the exclusion zone and that were innervated by the island or non-island RGC boutons. We posited that, in the absence of preference, each dendrite emerging from the cell body would have an independent probability for connecting to either island or non-island RGC boutons. We therefore counted the number of dendrites that emerged from each TC cell body that eventually connected to an RGC input (2,2,3,4,4,4,5,5,7). We then ran 100,000 trials in which each primary dendrite was assigned island or non-island connectivity with equal probability. For each iteration we counted the number of TCs that were innervated either only by island boutons or only by non-island boutons (captured). We found that 99% of the Monte Carlo iterations produced 4 (44%) or fewer captured TCs (Figure 2I). The 9 captured TCs (100%), therefore, represents a substantial deviation from an unbiased assignment of dendrites to island or non-island RGCs.

The absences of TCs that were innervated both by the island and non-island RGC boutons means that the island represents a segregation of retinogeniculate synaptic connectivity. This exclusivity is what we had predicted based on the model in which activity dependent synaptic remodeling would prevent strong functional dissimilarity in the RGC innervation pattern of TCs. We were surprised, though, at the extent of dendritic arbor asymmetry we observed associated with this connectivity pattern. Asymmetric thalamocortical cell arbors, particularly in the dorsal shell of the dLGN is not surprising (Krahe et al. 2011). What was noteworthy, was that the asymmetry was so clearly oriented away from the exclusion zone.

Few thalamocortical cell dendrites crossed into the exclusion zone and those that did, assumed a distal dendrite morphology (Figure 2H,J, Supplementary Figure 2,3, Supplementary Video). While most of these distal morphology neurites were left untraced, more complete tracings of two TCs confirmed that these dendrites maintained their non-RGC-receiving character when they passed out of the exclusion zone (TC3, TC4, Figure 2J, Supplementary Figure 2,3). The presence of cortical feedback synapses (small round boutons innervating small spines, (Guillery 1971)) on the dendrites that span the exclusion zone means that the cortical feedback onto the TCs does not obey the strict island/non-island segregation exhibited by the RGC innervation.

### Local inhibitory neurons follow RGC segregation

In mouse, RGC inputs to TCs are almost always coupled with connections to LINs (J. L. Morgan et al. 2016). These inhibitory neurons form three types of neurites: 1) Input/output shaft dendrites that span hundreds of micrometers and multiple subregions of the dLGN, 2) Input/output targeted dendrites that extend about 20 μm from the shaft dendrites and closely follow nearby RGC axons, and 3) Output-only axons that are relatively small (Josh L. Morgan and Lichtman 2020). The short distances between the output synapses of the targeted neurites and multiple RGC inputs means the targeted neurites can provide a synaptic drive that reflects the functional properties of the local dLGN neuropil (Josh L. Morgan and Lichtman 2020). Output synapses on the shaft dendrites of LINs, in contrast, are expected to be driven more by the global integration of inputs across the large LIN dendritic arbor. Based on these properties, we hypothesized that the shaft neurites of local inhibitory neurons would be indifferent to the boundaries of the RGC-island while individual targeted neurites would be associated with either island or non-island RGC boutons. We tested this hypothesis with both light and EM reconstructions of local inhibitory neurons.

In GAD67-GFP transgenic mice, dLGN local inhibitory neurons are brightly labeled with GFP (Seabrook et al. 2013; Charalambakis et al. 2019). We bred these mice with the albino tyr-/- mice to generate GAD67-GFP, tyr-/- transgenic mice (Figure 3A). Airyscan confocal images of the island revealed LIN dendrites that were closely associated with the CtB labeling of RGC boutons (Figure 3B). This tight association with RGC boutons is characteristic of the targeted dendrites of LINS. In contrast, the shaft dendrites of LINs crossed the RGC bouton exclusion zone without obvious deviation (Figure 3A,C). This pattern was consistent across five mice examined.

**Figure 3.**
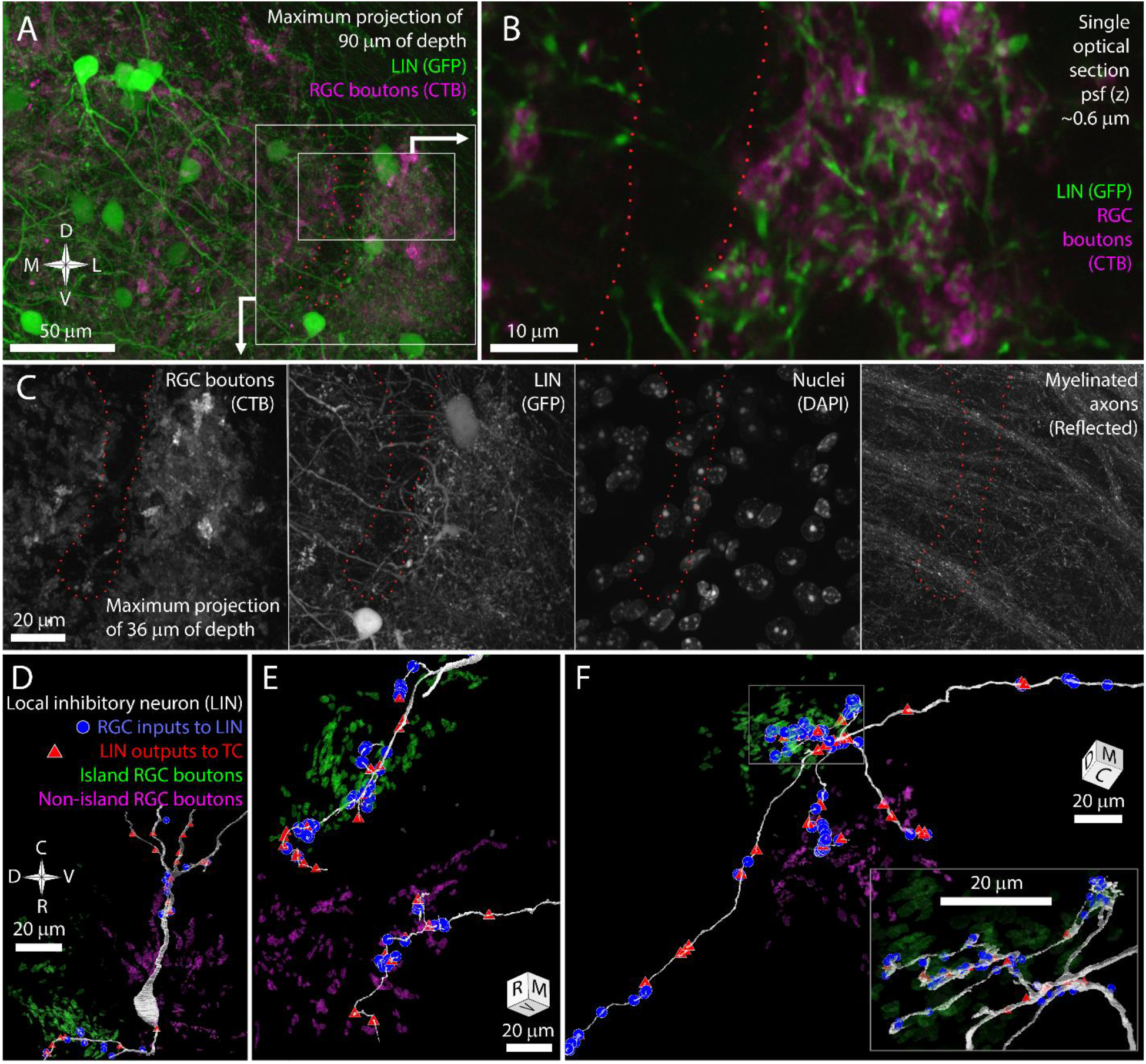
The long shaft dendrites of LINs cross the exclusion zone while the individual targeted dendrites of LINs have distict island or non-island synaptic identities. A) Reference image for panel B and C showing LINs (green) and CtB labeling of RGC terminals (magenta). B) Close association between targeted neurites of LINs (green) and RGC terminals (magenta). C) Four channels of confocal stack showing RGC bouton exclusion zone relative to LINs, cell nuclei, and reflective fibers. D) EM reconstruction of LIN relative to island (green) and non-island (magenta) RGC boutons. Input synapses from RGCs (blue circles) and output synapses (red triangles) are labeled. E) Two LIN dendrites labeled as in D. F) LIN dendrite labeled as in D. Inset shows magnified view of branch points. The functional identities of LINs are, therefore, likely to be locally, but not globally, specific to island or non-island boutons.

We next used our vEM volume to determine if the impression of shaft dendrite indifference to island boundaries and targeted dendrite avoidance of island boundaries was reflected in synaptic connectivity. We first selected one LIN for reconstruction based on the presence of its nucleus in the exclusion zone. This LIN extended dendrites into both the island and non-island neuropil and formed synapses with RGCs and TCs in both regions (Figure 3D). We next selected three LIN dendrites for tracing based on their synaptic connectivity with identified TCs and RGC boutons. Two of these tracings revealed long targeted neurites that formed many connections to TCs and RGCs specifically within either the island or non-island neuropil (Figure 3E).

In the third cell, we found seven dendritic processes extending from a small region near the interface of the island and the exclusion zone (Figure 3F). Four of the dendrites crossed the exclusion zone, formed no synapses with RGCs or TCs in this region, and then formed synapses with RGCs and TCs when they entered the non-island neuropil. One of these dendrites exhibited the thin diameter, tortuous path, and dense glomerular innervation of a LIN targeted dendrite. This dendrite was interesting in that it received four island boutons near its base and then received forty-two synapses from non-island RGC boutons.

The LIN results are consistent with our prediction that shaft dendrites would be indifferent to island/non-island boundaries while individual targeted dendrites would target either the island or non-island RGC boutons. However, the restriction of the targeted dendrites to one or the other RGC field does not appear to be an absolute rule. Rather the scale of targeted dendrite exploration and the size of the exclusion zone is likely to reduce the chances that a targeted dendrite would find partners on both in the island and outside of the island. This matching between the exploration of targeted LIN dendrites and the segregation of retinogeniculate connectivity means that targeted LIN dendrites will have an RGC input profile (island/non-island) that matches the TCs they innervate.

## DISCUSSION

The key conclusion of this study is that the previously reported islands of mistargeted RGC terminals represent a segregation of retinogeniculate circuitry into distinct pathways. The extent of this segregation was surprising. In a previous EM reconstruction of a wildtype mouse dLGN, nearby thalamocortical cells shared input from the same RGC axons even when the RGC axons or TCs were of morphologically distinct types (J. L. Morgan et al. 2016). Based on these results, we argued that the parallel pathways of different types of RGCs did not have synaptically clean boundaries in the mouse dLGN, but instead were frequently intermixed. In contrast, the island / non-island projections appear truly parallel (non-intersecting). The fact that there is a developmental starting condition that can induce a synaptically segregated microcircuit has important implications for our understanding of the organization of visual circuits and for our understanding of the implementation of activity dependent development.

### The network organization of mouse dLGN

In cats and primates, the receptive field properties of TCs are usually derived from strong innervation from a few functionally similar RGCs (Usrey, Reppas, and Reid 1998; Mastronarde 1992; Hamos et al. 1987). This connectivity allows for a spatially continuous sampling of visual space with little loss in acuity (Alonso et al. 2006; Martinez et al. 2014). Studies of retinogeniculate connectivity in mouse suggested a messier view of dLGN processing. Mouse TCs can be innervated by dozens of RGCs (J. L. Morgan et al. 2016; Hammer et al. 2015; Rompani et al. 2017), and different RGC types innervate the same TC (L. Liang et al. 2018; Marshel et al. 2012; Rompani et al. 2017). Part of the explanation of the discrepancy is that the mouse dLGN contains both high and low-convergence TCs (J. L. Morgan et al. 2016; Rompani et al. 2017). Further, many of the RGCs inputs to a given mouse TC are much weaker than the relatively few RGCs that dominate the TCs firing (Litvina et al. 2018). Within the circuitry of the mouse dLGN, therefore, there are TCs whose connectivity is consistent with the gated relays of retinal activity described in other species.

However, the most striking connectivity rule that came out of our mouse EM reconstruction was that we could find no distinction between TCs (morphology, convergence, ultrastructure) that precluded two TCs from being innervated by the same RGC axon (J. L. Morgan et al. 2016). We wondered if the seeming lack of synaptic specificity was because mouse evolution valued signal detection over functional specificity, if the mixing of channels was a statistical selection of related channels, or if synaptic specificity was limited by the size of the tissue. A whole mouse dLGN can fit inside one of the six functionally distinct sublamina of a macaque dLGN (Casagrande et al. 2007). The odd case of the albino island eliminates the possibility that the mixing of retinogeniculate connections observed in the normal mouse reflects an upper limit of the structural specificity of the system. The mechanisms exist in the mouse dLGN to generate completely segregated subcircuits. The normal network organization, therefore, likely reflects an adaptive balance between receptive field refinement, space, and the benefits of integrating signals from multiple RGCs.

### Evidence for activity dependent remodeling as the driving force behind islands

How much of the circuit segregation of the albino dLGN islands can we attribute to activity dependent remodeling? There are activity independent mechanisms that could be contributing to the island phenotype. Molecular recognition helps keep RGC axons from similar regions of the dLGN bundled together as they travel through the optic tract (Sitko, Kuwajima, and Mason 2018; Bruce et al. 2017). Homotypic membrane recognition molecules are also required for RGC boutons to cluster together to form synaptic glomeruli (Monavarfeshani et al. 2018). A set of mistargeted RGC axons of the same type might, therefore, be expected to stay bundled together in the absence of activity dependent cues.

On the other hand, there is extensive evidence showing that developmental patterns of retinal activity are critical for shaping the architecture of retinogeniculate connections (Liang Liang and Chen 2020 for review). The waves of activity that propagate through the retina prior to eye opening are particularly important for eye-specific segregation (Feller 2009) and topographic refinement(Grubb et al. 2003; Pfeiffenberger, Yamada, and Feldheim 2006). Blocking the propagation of these waves also prevents the formation of RGC bouton islands in albino mice and EphB1 KO mice (Rebsam, Bhansali, and Mason 2012; Rebsam, Petros, and Mason 2009). Thus, the segregation of island and non-island RGC boutons depends on the same activity dependent synaptic remodeling process that is responsible for refining the receptive fields of TCs in the normal mouse dLGN.

The connectivity of the RGC islands reported here also support the conclusion that activity dependent circuit remodeling generates the islands. First, the island is not a neuroma of tangled RGC axons, and the exclusion zone is not a region of pathological dLGN. Rather, the island and exclusion zone represent a segregation or retinogeniculate connectivity within an otherwise normal dLGN neuropil. While activity independent signals might drive dissimilar RGC boutons apart it doesn’t explain why a TC could not be innervated by both island and non-island RGC boutons. The simplest explanation for clustering of RGC boutons, the binary connectivity of TCs, and the systematic asymmetry of the TC dendritic arbors is that the island is the result of activity dependent Hebbian synaptic remodeling.

### A model of activity dependent synaptic remodeling

The process of activity dependent synaptic remodeling begins with RGC axons having already positioned themselves in roughly the correct region of the dLGN (Godement, Salaün, and Imbert 1984). However, each TC receives input from many more RGCs than will eventually drive it (Chen and Regehr 2000) and its receptive field properties are likewise less refined than they will be in the adult (Tavazoie and Reid 2000; Tschetter et al. 2018). The dLGN then undergoes a period of activity dependent remodeling in which spatio/temporally structured retinal activity is thought to provide the information TCs need in order to select which retinal inputs to maintain and which to eliminate (Hong and Chen 2011).

Unlike many of the model systems where synapse elimination has been well characterized (Redfern 1970; Crepel, Mariani, and Delhaye-Bouchaud 1976; Jeff W. Lichtman 1977), the synaptic remodeling of TCs is not all or none. Functionally similar RGCs can maintain and elaborate their connections on the same TC. The capacity of functionally similar synapses from different axons to reinforce one another may result from the fact that the activity dependent synaptic enhancement of retinogeniculate synapses acts on the time scale of bursts of activity (∼1s) instead of individual spikes (Butts, Kanold, and Shatz 2007). A mature RGC can, therefore, share a TC target with a slightly more ventral RGC which shares a target with a even more ventral RGC and so on in every direction. The result of dLGN remodeling is a refined but also synaptically continuous remapping of visual space.

In albino mice, where sets of RGCs make large and correlated errors in the initial targeting of retinogeniculate projections, synaptic stabilization is no longer equal in every direction. A group targeting error produces sets of axons whose firing patterns are strongly correlated within the group but that are different from the firing patterns of surrounding RGC axons. At the start of synaptogenesis, the dLGN neuropil at the boundary between the mistargeted and normally targeted RGC axons would be composed of TC dendrites whose RGC inputs are systematically less correlated than dendrites oriented away from the boundary. Mistargeted RGCs would be more likely to stabilize synapses in the center of the field innervated by the mistargeted RGCs. This process would also be self-reinforcing. As boundary synapses are weakened in favor of centrally located synapses, centrally located TC targets become even more attractive than boundary targets. The mechanism driving mistargeted retinogeniculate connections to collapse into a completely segregated island would not be different from the mechanisms of normal activity dependent development. The difference is simply that in the normal dLGN, synaptic stabilization forces are much closer to being equal in all directions.

### Why is there an exclusion zone?

Hebbian synaptic remodeling predicts mistargeted axons will capture a set of unique TC targets, but it does not predict the spatial segregation of synapses we observe in the form of the exclusion zone. The cell bodies of island and non-island innervated TCs are close enough together that their RGC-bouton- receiving dendrites should overlap. If the only remodeling rule implemented was to prune the weaker population of synapses on TCs innervated by both island and non-island RGCs, then we would expect to find the spared RGC boutons filling the boundary region between the mistargeted and normally targeted RGC axons. The exclusion zone requires an additional level of circuit remodeling.

An explanation for the exclusion zone is suggested by the asymmetry of the TC dendritic arbors relative to the exclusion zone. The asymmetry would seem to require that developing TCs direct dendritic resources towards favored inputs. TC dendrites normally respond to RGC inputs by extending spines around input RGC boutons (Wilson, Friedlander, and Sherman 1984). If TCs are deprived of retinal inputs during development, they undergo a brief period of exuberant dendritic growth followed by dendritic pruning (El-Danaf et al. 2015). In adult dLGN, loss of RGC inputs leads to an eventual shrinkage of dendritic arbors (Bhandari et al. 2022). If the trophic interaction between RGC boutons and TC dendrites can operate with dendritic specificity, then the exclusion zone could represent a region where dendrites innervated by functionally disparate RGC inputs were pruned in favor of dendrites in regions where RGC inputs were correlated.

## CONTRIBUTIONS

SM, XC, and JH injected eyes and collected tissue. SM, XC, JH, and JM collected optical images. KV processed tissue for EM. LM and JM collected EM images. JM wrote the analysis/rendering code. JM and SM planned experiments. JM and PW provided funding. JM wrote the manuscript. All authors read and approved the manuscript.

## ACKNOWLEDGEMENTS

Thanks go to the Washington University Center for Cellular Imaging and staff where the EM was performed and to ScalableMinds that aligned the EM dataset. Thanks to Richard Schaleck and the Jeff Lichtman lab providing carbon coated Kapton collection tape. This work was supported by an unrestricted grant to the Department of Ophthalmology and Visual Sciences from Research to Prevent Blindness, by a Research to Prevent Blindness Career Development Award (J.L.M), and by the NIH (EY029313 to J.L.M).

## ETHICS DECLARATIONS

The authors have no conflict of interest to declare.

---

**Supplemental Figure 1.**
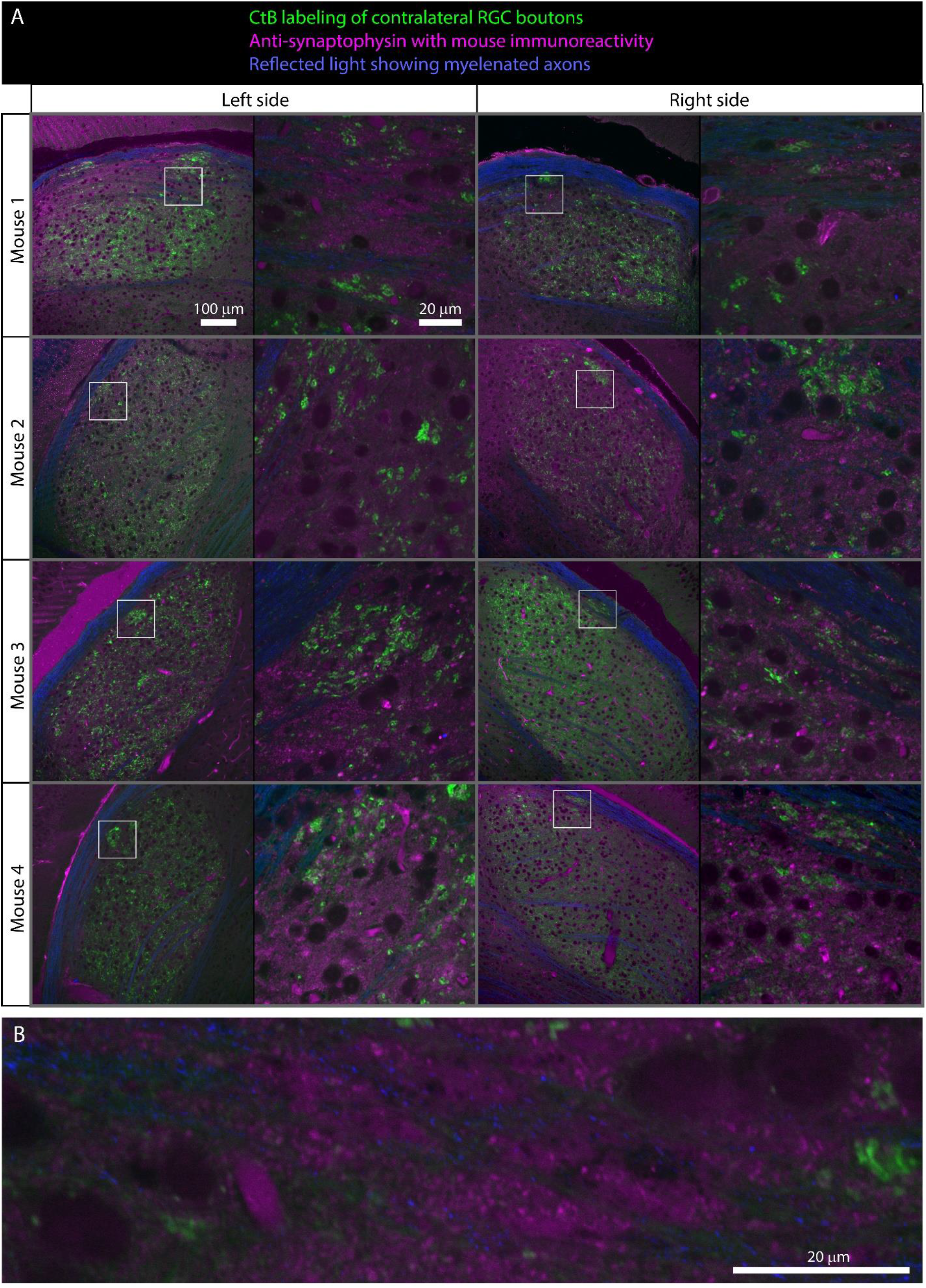
Synaptophysin labeling of dLGN synapses shows synaptic neuropil is present in the RGC bouton exclusion zone. A) Each row show the left and right dLGN is shown for each mouse. Contralateral projecting RGC terminals labeled with CtB (488 or 555) are shown in green. Mouse anti-synaptophysin immunoreactivity is shown in magenta. Anti-mouse secondary antibody labels blood vessels as well as anti-synaptophysin primary antibody. Reflected light highlighting myelinated axons is shown in blue. A) For each dLGN, as single plane of a confocal scan of the full dLGN (left) and exclusion zone (right) is shown. The white box indicates the position of the high-resolution image relative to the full dLGN. Bright punctate immune artifacts are visible in some panels. B) Closer look at exclusion zone of left dLGN from mouse 1 shown in panel A. The immunolabeling showing synaptic neuropil in the exclusion zone is consistent with our examination of the EM volume.

**Supplementary Figure 2.**
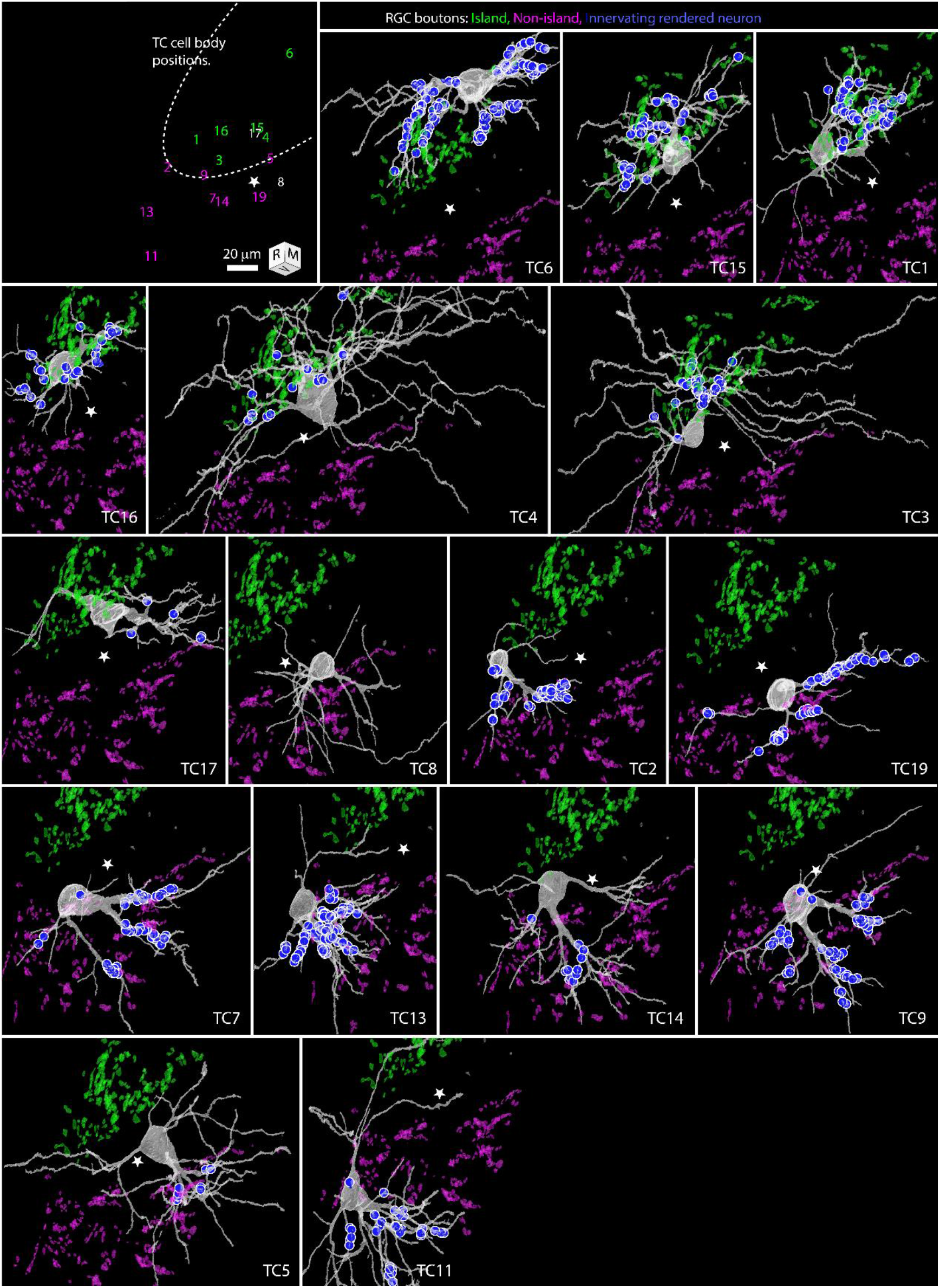
TCs surrounding the exclusion zone are captured by either the island or non-island RGC boutons. The TC dendritic arbors reflect this capture. Panels show renderings of partially reconstructed TCs. The top left key shows the position of the nuclei of the reconstructed TCs relative to the boundary of the RGC island (dotted white line). The color of the cell IDs indicates whether they received RGC input from the island or non-island RGC boutons. The star indicates a common reference position for all panels. RGC inputs innervating the TC highlighted in each panel are shown as blue circles. The TC connectivity and morphologies are consistent with Hebbian rules shaping retinogeniculate circuit structure.

**Supplementary Figure 3.**
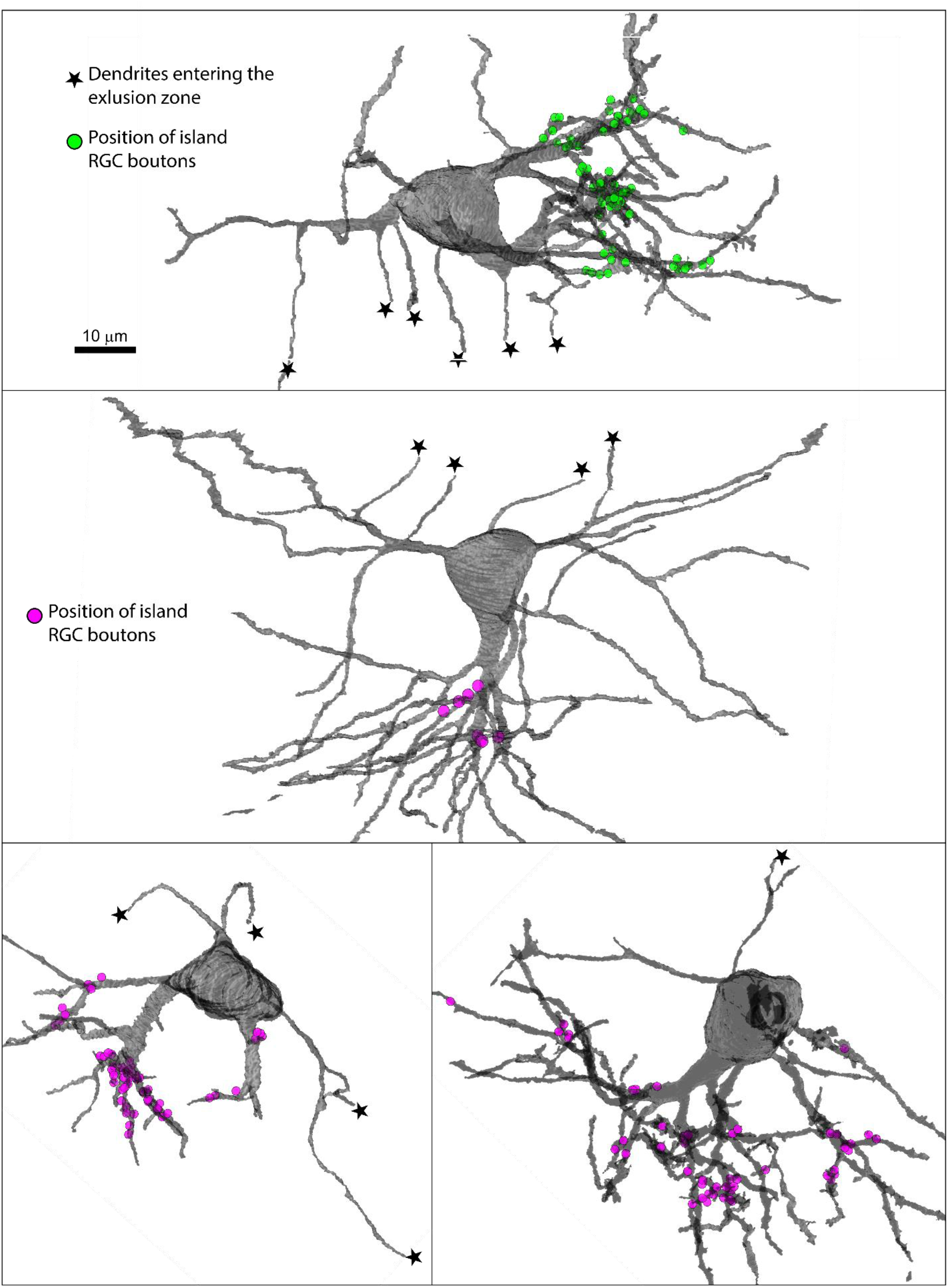
Proximal dendrites of TCs are usually thick and innervated by RGC boutons. Proximal dendrites of TCs that extend into the exclusion zone are noticeably thinner, similar to distal dendrites. Panels show four example TCs where dendrites were observed extending into the exclusion zone. Neurites in the exclusion zone are indicated with stars. The location of RGC boutons innervating the TC are indicated by circles (green = island, magenta = non-island).

## Notes

### Competing Interest Statement

The authors have declared no competing interest.

### Summary of Updates

We have made minor changes to improve clarity.

