## Supplementary figures and images for "Mistargeted retinal axons induce a synaptically independent subcircuit in the visual thalamus of albino mice"

### Supplemental Figure 1

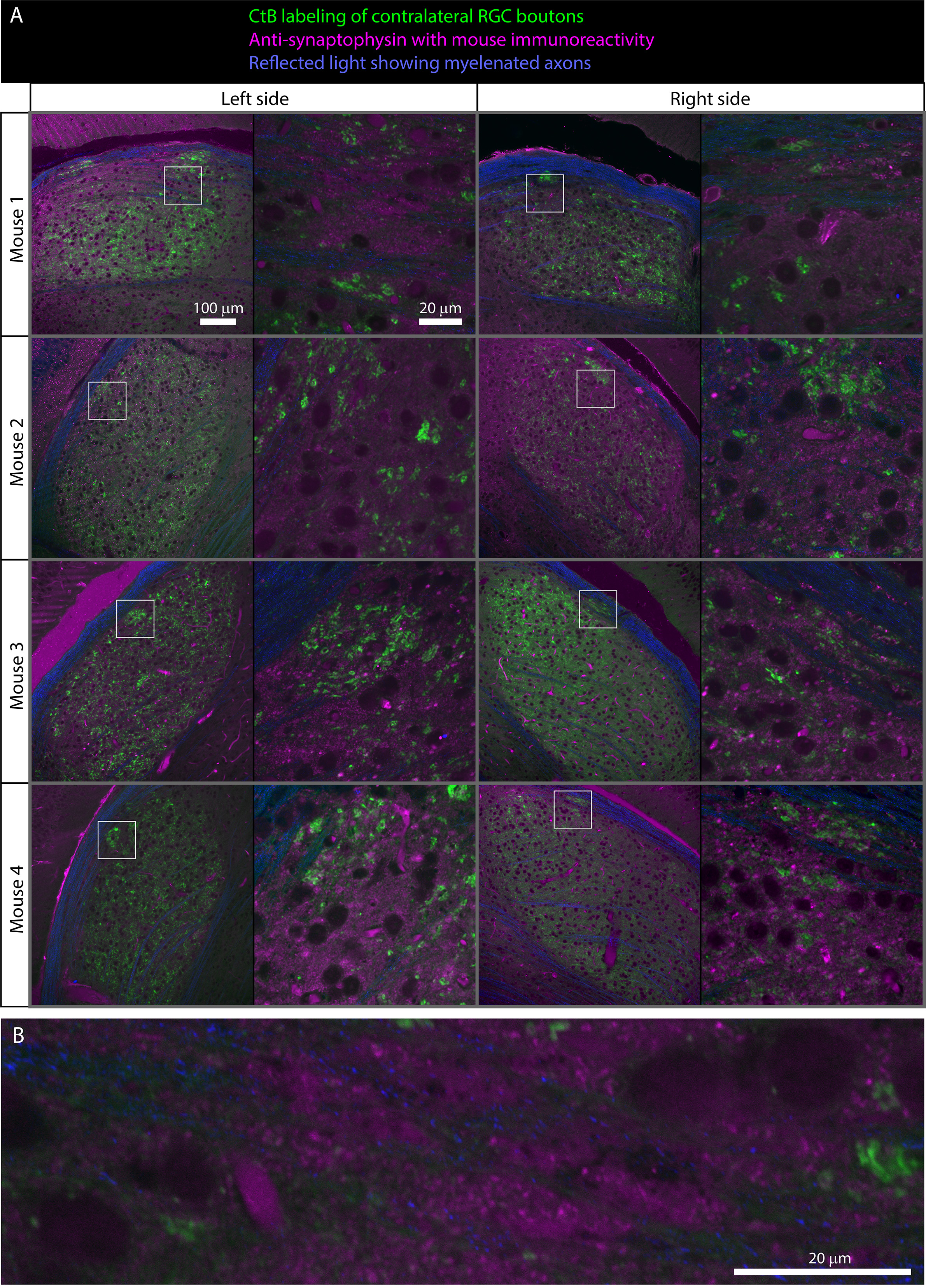

### Supplemental Figure 2

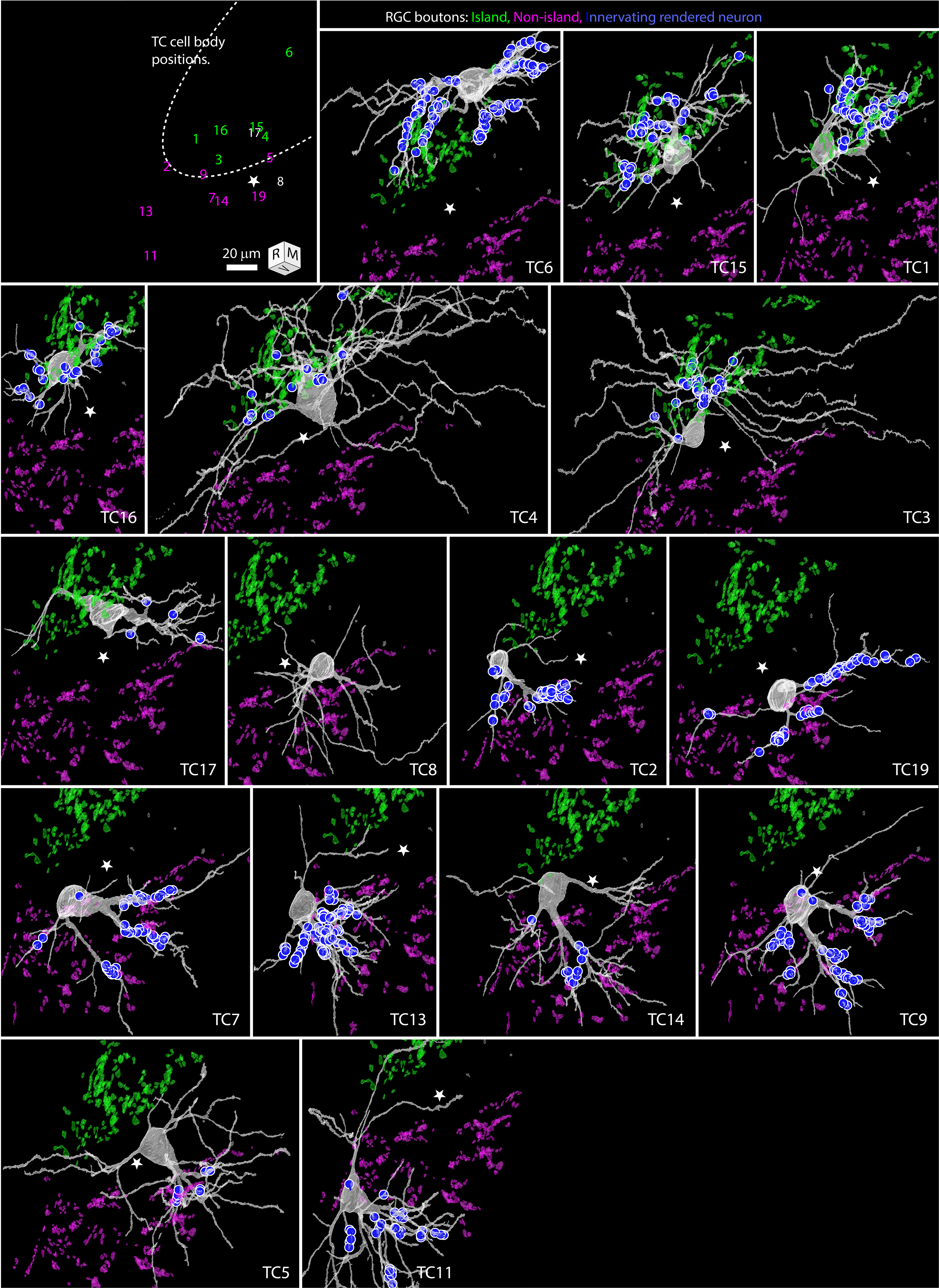

### Supplemental Figure 3

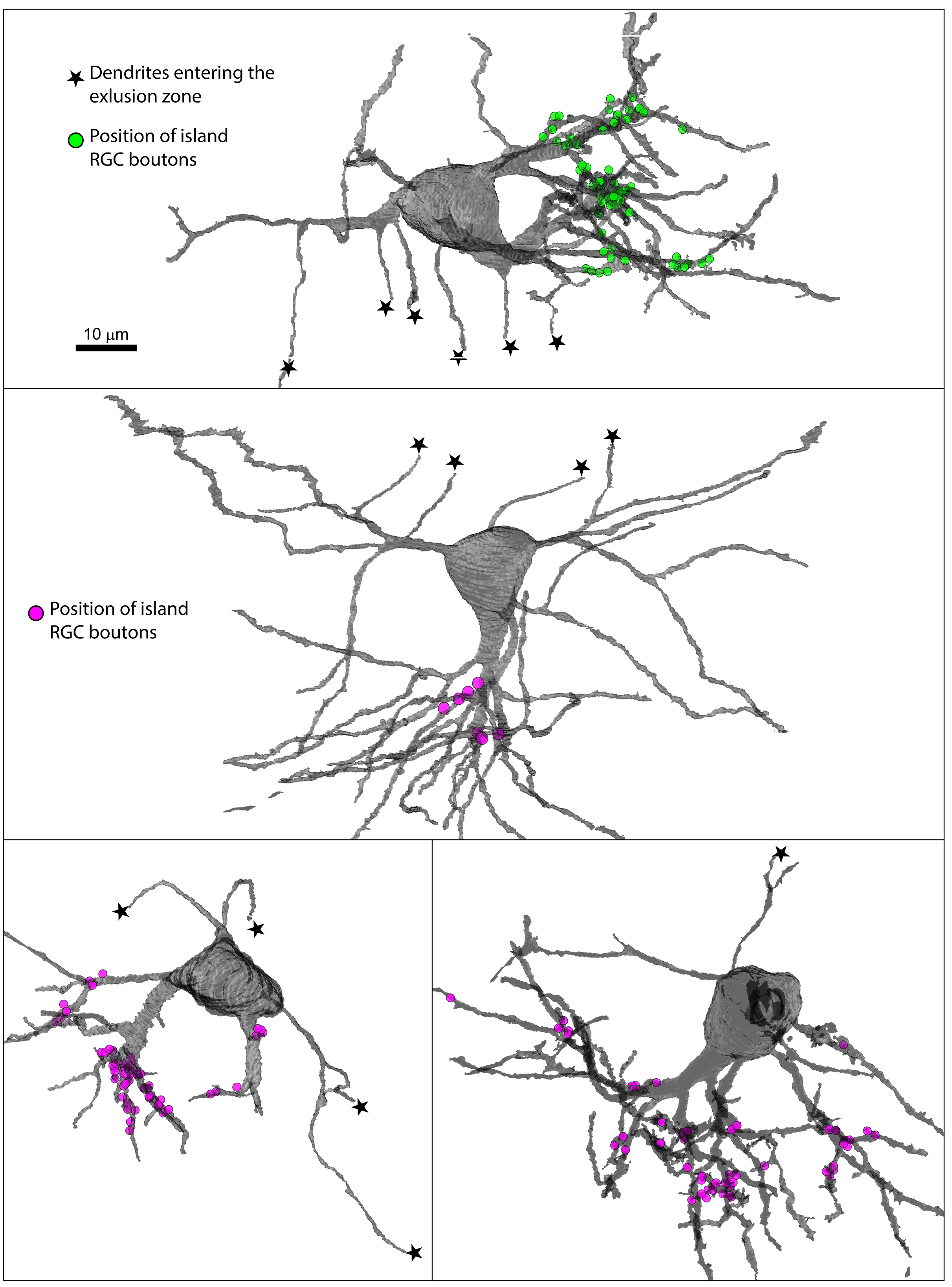
